# Food Webs Over Time: Evaluating the Variability of Degree Distribution on Ecological Networks

**DOI:** 10.1101/134833

**Authors:** Daniela N Lopez, Patricio A Camus, Nelson Valdivia, Sergio A Estay

**Author notes:** Corresponding author: Daniela N. López, Instituto de Ciencias Ambientales y Evolutivas, Facultad de Ciencias, Universidad Austral de Chile. Edificio Emilio Pugin, tercer piso oficina 313,Valdivia, Chile. Telephone number: 056+ 63+ 2221344, Correspondence:.

## Abstract

Although networks analysis has moved from static to dynamic, ecological networks are still analyzed as time-aggregated units where time-specific interactions are aggregated into one single network. As a result, several questions arise such as what is the functional form of and how variable is the topology of time-specific versus time-aggregated ecological networks? Furthermore, it is yet unknown to what extent the structure of time-aggregated networks is representative of the dynamics of the community. Here, we compared the topology of time-specific and time-aggregated networks by analyzing a set of intertidal networks containing more than 1,000 interactions, and assessed the spatiotemporal dynamics of their degree distributions. By fitting different distribution models, we found that the out-degree distributions of seasonal and time-aggregated networks were best described by an exponential model while the in-degree distributions were best described by a discrete generalized beta model. The degree distributions of the seasonal networks were highly temporally variable and are significantly different from those of time-aggregated networks. We observed that seasonal degree distributions converged toward time-aggregated network distributions after 1.5 years of sampling. Our results highlight the importance of understanding the dynamics of ecological networks, which can show topological characteristics significantly different from those of time-aggregated networks.

## Introduction

In the last decades, the study of networks has moved from static to dynamic analyses; thus, deeper insights into the internal structure of networks at different temporal and spatial resolutions can be obtained ^1^. Despite this, trophic network analyses are often based on time-aggregated networks constructed by combining all interactions observed during a wide time frame. In time-aggregated networks, interactions occurring at different periods of time, for example during different seasons of the year, are summarized into a single network. Although such networks are powerful abstractions ^2^ that have proven to be fruitful in yielding insights into overall network properties, they fail to represent the range of dynamics occurring in a system ^3^. Further to this, in some cases these static representations have been shown to yield misleading results regarding the importance of species or interactions to the persistence of the network ^4,5^.

The analysis of dynamic trophic networks has been addressed from empirical ^4,6^ and theoretical ^6^ perspectives to understand how changes in network topology modify the flow of energy through the network. Due to the inherent dynamic nature of natural communities, it is expected that the architecture of trophic interactions should be highly variable ^4^. For example, one of the few studies that incorporates temporal variability into a trophic network shows that topological characteristics can vary significantly between different periods of time ^2^. In this sense, determining the temporal scale that captures relevant patterns in the structure and persistence of networks is highly relevant.

An approach to the study of dynamics of network topology is through the analysis of the degree distribution. In the case of trophic networks, the degree distribution is among the most important topological descriptors as it allows for the identification of important interactors ^7^ and describes general patterns of energy flux across the network ^8^. Several studies have shown that the degree distribution of trophic networks has various functional forms, and most examples deviate significantly from Erdos-Reyi networks ^9^,^10^, ^11^, ^12^. Also, given the natural direction given by the flux of energy from primary producers to predators ^13^ a clear difference between in and out-degree distribution exists. In trophic networks the in-degree distribution represents the number of prey consumed by each predator (niche breadth), while the out-degree represents the number of predators attacking each prey ^14, 15,16^In this vein, given that the in- and out-degree distributions depict two different ecological processes, we would expect that their distributions would have different functional forms.

By explicitly considering the temporal dynamics of interactions, temperate intertidal rocky-shore communities provide an opportunity to compare time-aggregated networks with time-specific networks. A recent study by López et al. ^17^ shows that northern Chilean rocky shore communities can harbor a large number of transient trophic interactions (≈80% of ca. 1,000 interactions), which might indicate that the degree distribution of these networks is seasonally dependent. To contribute to the understanding of trophic network dynamics, in this study we analyzed the spatiotemporal dynamics of the degree distribution of a set of large intertidal rocky-shore trophic networks sampled at high spatial, temporal, and taxonomic resolution. Throughout the study we address the following questions: a) What is the functional form of and how variable are the in- and out-degree distributions? b) What is the relationship, if any, between the topological characteristics of seasonal (time-specific) and time-aggregated networks? c) What is the temporal scale at which topological descriptors converge to their aggregated values? And finally, d) if the characteristics of time-aggregated networks diverge from those of seasonal networks, then what do the former tell us about natural communities?

## Results

### Degree distribution

We observed a strong decay in the rank-frequency of the degree distributions for all networks (Fig. 1). While the in-degree distribution was best fit to the DGBD function (Table 1), the out-degree distribution was better represented by the exponential distribution (Table 1). Furthermore, the in- and out-degree distributions were highly temporally variable (Fig. 2, Table 1) as seen in the values of the parameters (*a*, *μ*) of the fitted seasonal distributions (Table 1). The site-specific estimated values of the *a* parameter of the DGBD for the in-degree distributions were (minimum, maximum, and time-aggregated networks): RS (0.017; 0.54; 0.18), CC (0.037; 0.67; 0.058), CA (0.02; 0.40; 0.054) and LA (0.061; 0.415; 0.247). Regarding the out-degree distribution, the values of the *a* parameter were: RS (0.55; 0.81; 0.62), CC (0.65; 0.85; 0.67), CA (0.85; 0.91; 0.87) and LA (0.84; 0.92; 0.88) (Fig. 2, Table 1). The values of the parameters of the exponential distribution (*μ*) (minimum, maximum, and time-aggregated networks) for the in-degree distribution were: RS (0.05; 0.13; 0.03), CC (0.05; 0.12; 0.03), CA (0.05; 0.15; 0.03), LA (0.05; 0.13; 0.03) (Fig. 2, Table 1). In the case of the out-degree distribution, the estimated parameters of the exponential distribution were: RS (0.185; 0.41; 0.14), CC (0.185; 0.58; 0.13), CA (0.167; 0.56; 0.12), LA (0.161; 0.42; 0.12) (Fig.2, Table 1).

**Figure 1:**
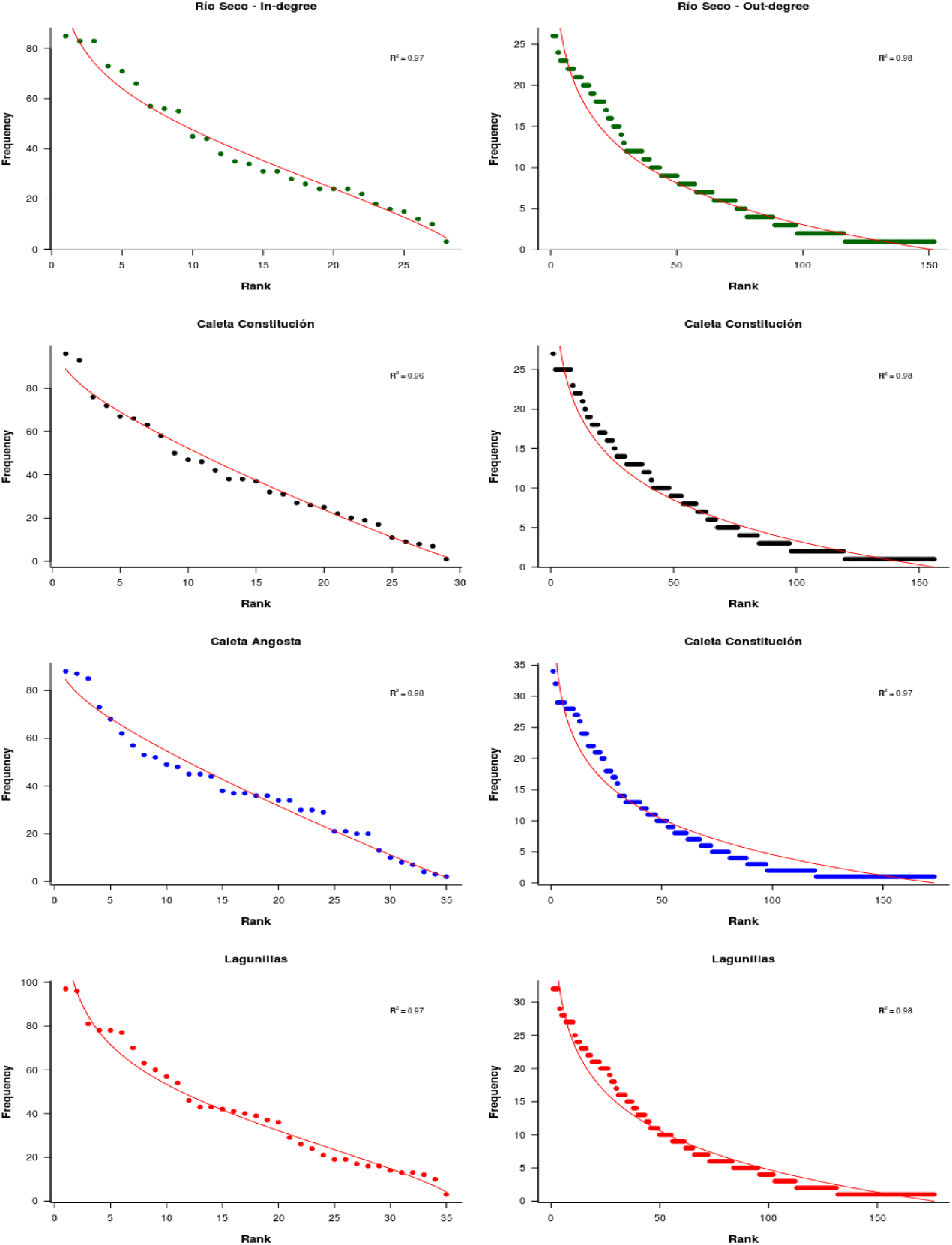
The rank-order of the in-degree distribution (left) and exponential distribution of the out-degree (right) distribution of time-aggregated intertidal networks of each locality of the study. Color codes are as follows: Río Seco (dark green), Caleta Constitución (black), Caleta Angosta (blue), Lagunillas (red).

**Table 1:**
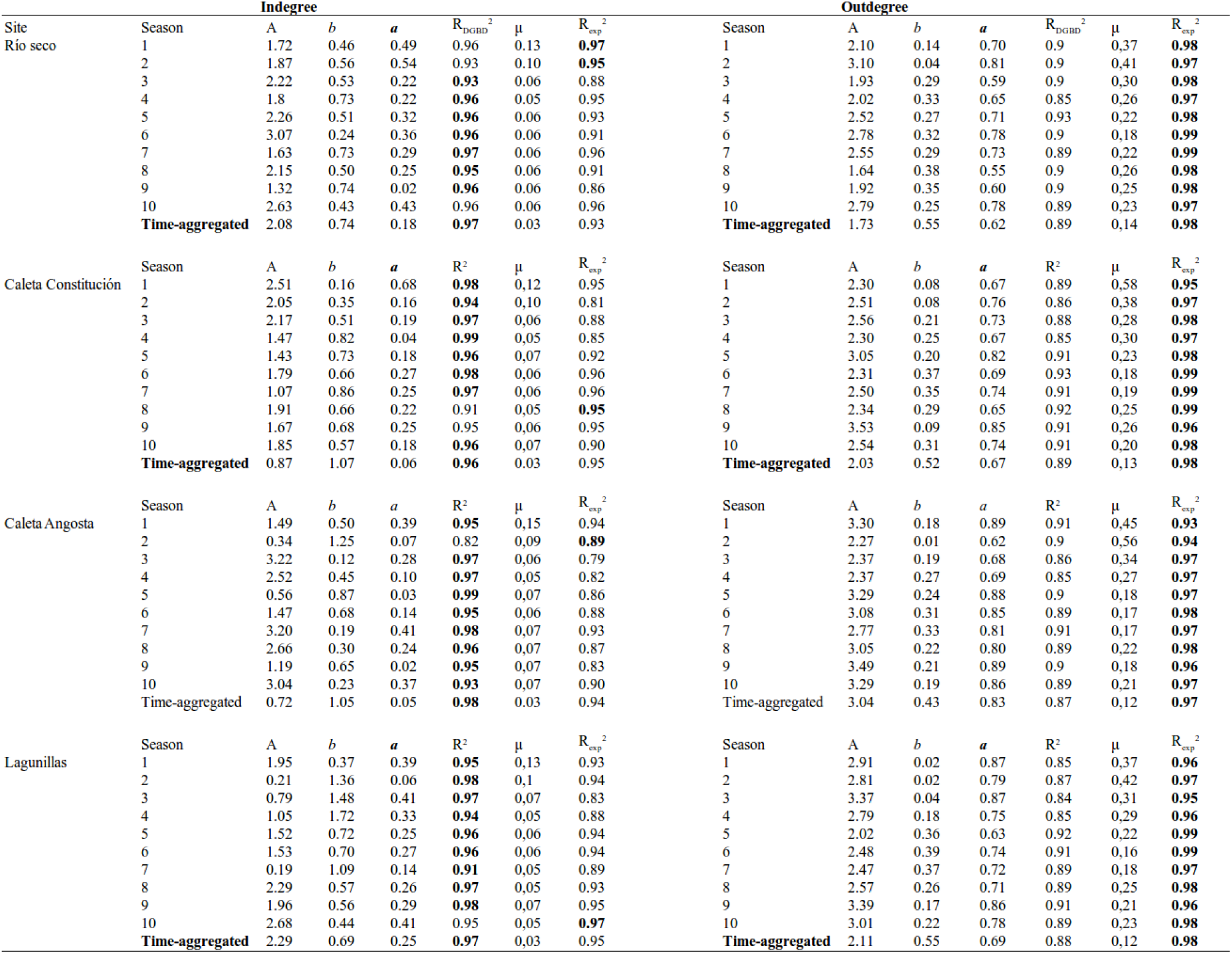
Parameters of the DGDB (A, *b, a*, R^2^) and exponential distribution (μ, R^2^) for the seasonal and time-aggregated networks for each locality of the study. Seasons range from spring 2004 to summer 2007.

**Figure 2:**
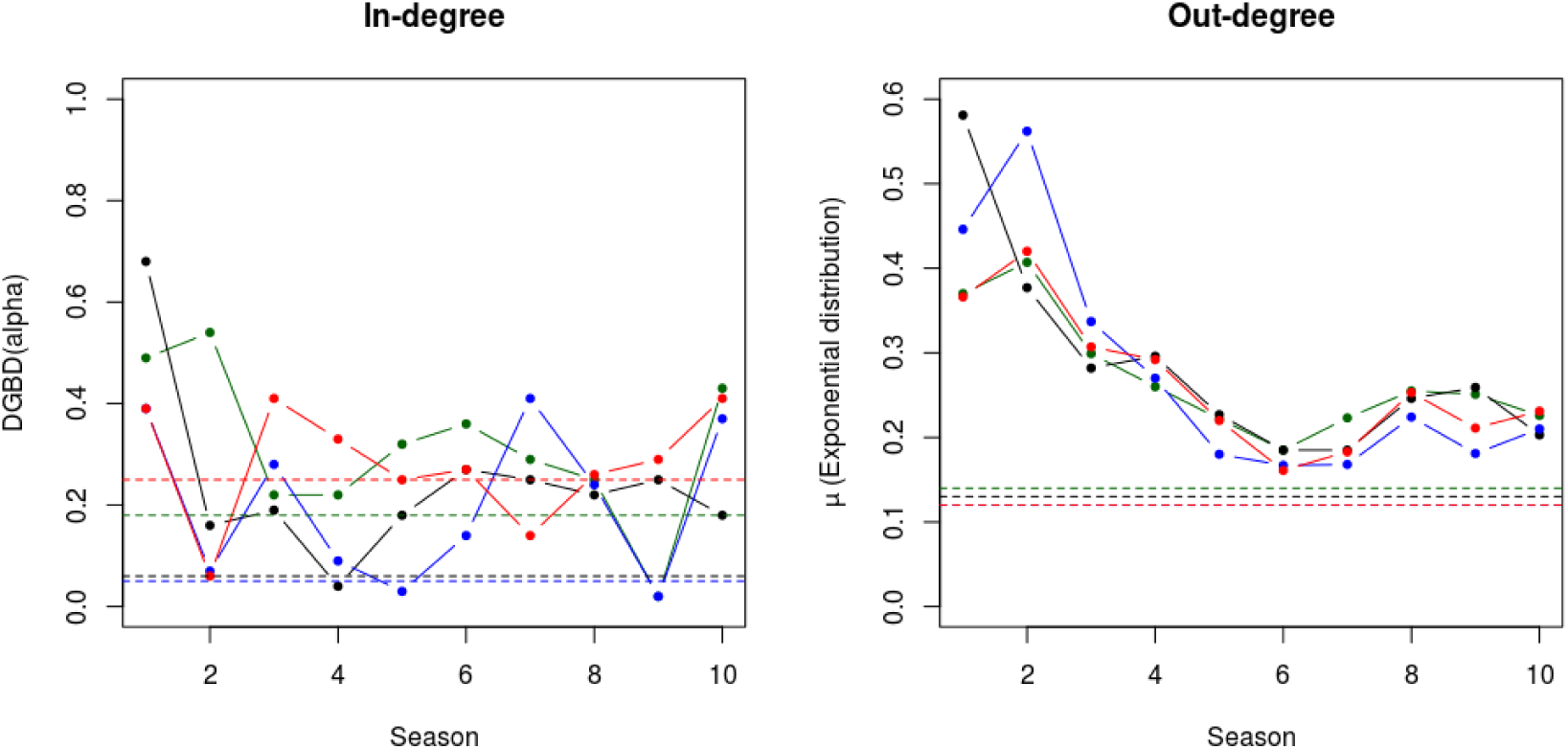
Temporal variability of *a* of the DGDB for the in-degree distributions (left) and μ of the exponential distribution for the out-degree distributions (right) for in each locality of the study. Color codes are as follows: Río Seco (dark green), Caleta Constitución (black), Caleta Angosta (blue), Lagunillas (red). Dotted lines indicate the estimated value for the time-aggregated networks.

### Convergence

Most of the *a*-values (DGBD) of seasonal network in-degree distributions were slightly higher than those of the time-aggregated networks (Fig. 2, Table 1). For the out-degree seasonal networks, values of *μ* in the exponential distribution were always above those of the time-aggregated networks (Fig. 2, Table 1). In the in-degree distribution, when we estimated *a* by sequentially adding seasonal networks, we found that convergence to the values observed in time-aggregated networks occurred between one and 1.5 years (four to six sampling events, Fig. 3). The same result was found for the sequential addition of the out-degree seasonal networks where the *μ* parameter converged to the time-aggregated values after roughly 1-1.5 years (Fig. 3).

**Figure 3:**
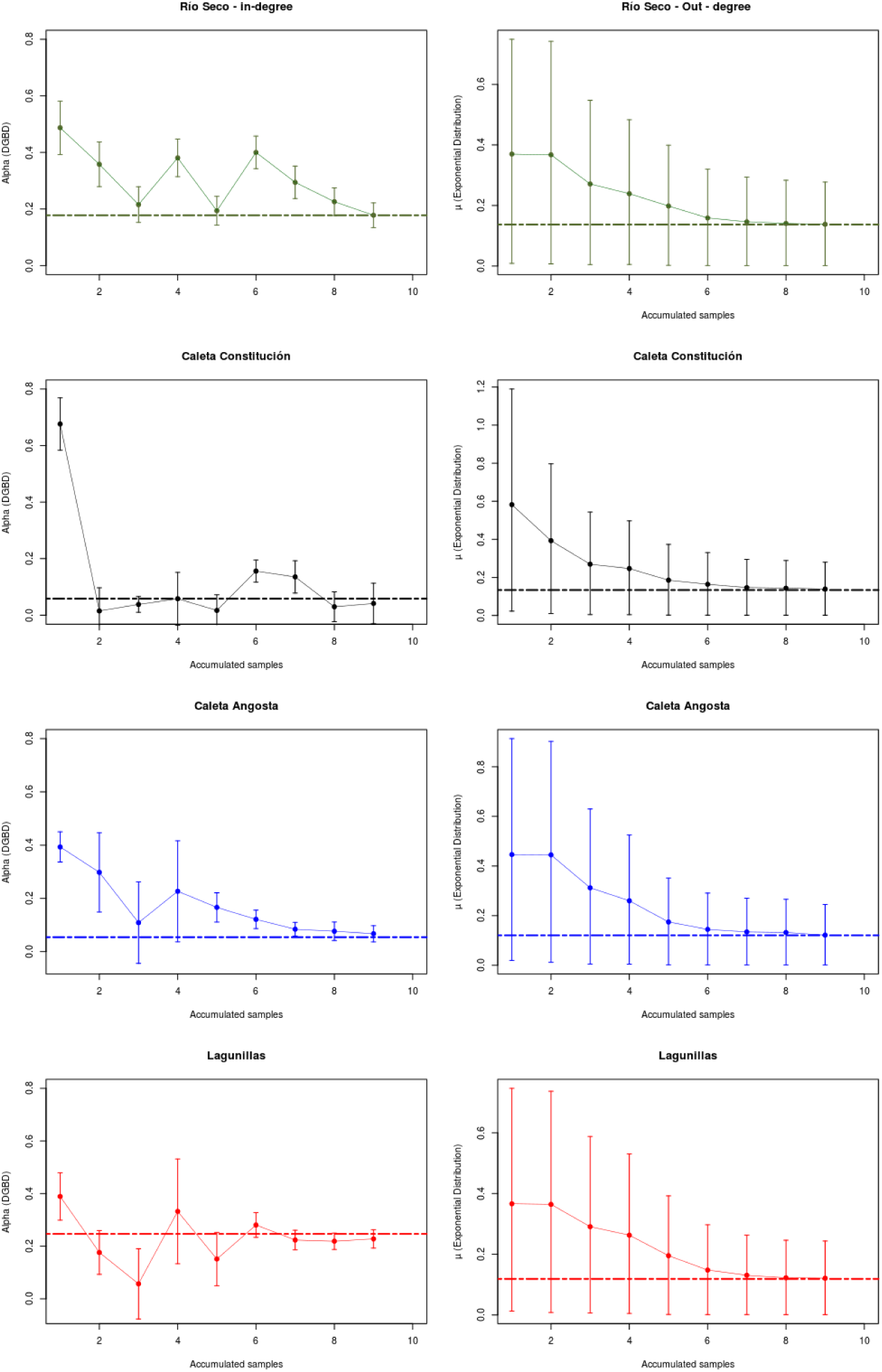
Convergence of *a* of the DGBD for in-degree distributions (left) and μ of the exponential distribution for the out-degree distributions (right) for each locality of the study: Río Seco (dark green), Caleta Constitución (black), Caleta Angosta (blue), Lagunillas (red). Dotted lines indicate the estimated value of the parameter for the time-aggregated networks.

## Discussion

We observed a common pattern of decay and a high temporal variability in the in and out-degree distributions of these intertidal rocky-shore assemblages. The best model for describing the in-degree distributions was a DGBD, and an exponential distribution best fit the out-degree distributions. These results indicate that the in- and out-degree distributions are generated by two different processes. The ecological significance of in-degree distributions is quite direct for trophic networks: they represent the distribution of trophic niche breadth across the species in the community ^18^. On the other hand, out-degree distributions portray predation pressure (number of predators consuming a type of prey) within a community; thus, it is difficult to relate the out-degree distribution with a niche axis. However, we can speculate that if some level of correlation exists between the number of predator species consuming a given type of prey and the total abundance of individual predators attacking it, then out-degree values represent at some point the enemy-free space available for the prey species ^19, 20^. Overall, however, deeper theoretical and empirical development is needed to assess the validity of this idea.

The results found here provide evidence that a given community can show a spectrum of topological characteristics over time. Other studies, in line with our results, have shown that network features such as niche breath and the number of interaction partners are highly spatially and temporally variable ^33^. In particular, the observed differences in the *a* and *μ* values between seasonal and time-aggregated degree distributions suggest that the isolated analysis of the latter can lead to misleading conclusions. The *a* and *μ* parameters represent how fast the degree distribution decays from hubs (most connected nodes) to one-degree nodes: the higher the value the faster is the decay. In trophic networks, if we sort in descending order the species rank according to its trophic niche breadth (in-degree) or the number of predators attacking it (out-degree), higher values of these parameters indicate larger differences between the most connected species. In our study, the fact that in most cases (>80%) the degree distribution of the seasonal networks had higher *a* and *μ* values than those of the time-aggregated networks means that hubs are more dominant (relatively much more connected ^34^) in the seasonal networks than in the time-aggregated networks. For example, in the case of the in-degree distributions, this means that the difference in terms of niche breath between the first- and second-rank species is several times larger than what is suggested by the analysis of the time-aggregated networks. The same is seen with the out-degree distributions: the species consumed for more predators is several times more connected than what is suggested by the analysis of the time-aggregated networks. A consequence of these discrepancies in the degree distributions of the seasonal and time-aggregated networks is a difference in the vulnerability of the networks to attacks. Due to its relatively higher dominance of hubs, seasonal networks are more vulnerable to the elimination of the most connected nodes than are the time-aggregated networks ^6, 7^. In ecological terms, the seasonal networks predict that the extinction of the most generalist predators or the most consumed species would have worst consequences for the community structure than that predicted using time-aggregated network analysis^21^. Despite this, in these particular networks, these potential negative effects could be counteracted by high functional redundancy, minimizing the impact of removing a hub ^22^.

The parameter values estimated from the DGBG and exponential distributions for the in- and out-degree distributions were highly asymmetrical (Table 1). A similar result was found in directed industrial networks where asymmetry between in- and out-degrees has been explained due to the fact that the in-degree distributions have sharp cut-offs and substantially lower degrees than those of the out-degree distributions ^23^. In our case, this indicates that, in trophic networks, the out-degree will always be larger than the in-degree given that the number of prey species is greater than the number of consumer species according to the pyramidal structure of these networks ^22 24^.These results encourage us to think on how are network structure and function related at biologically relevant time-scales? To resolve this question, networks including mechanistic information, such as functional responses and prey-switch behavior in the case of food webs, should be explored ^25^. No doubt the incorporation of this information will be a significant improvement to the conceptual framework of dynamic networks; however, until now such a task represents a mathematical challenge due to the complex analysis of multi-species assemblages ^24^.

Regarding temporal scales, our results showed a convergence of the *a* and *μ* parameter values of the seasonal networks to those of the time-aggregated networks after the sequential addition of four to six temporal networks. From a practical point of view, this means that, at least in this marine system, the structure of the whole community, including its interactions, will be captured only after sustained sampling effort; sampling at short time scales will yield gross misestimations of the statistical distribution of the in- and out-degrees in time-aggregated networks. Thus, our results illustrate that the description of a network depends on the time frame used in the monitoring scheme, and this could lead to different strategies of management for a given ecosystem ^2^. From a theoretical point of view, our results show that time-aggregated networks do not necessarily show similar topological characteristics than those of temporal networks ^4^. If the observed state of a community at a given time point differs from that predicted by a time-aggregated network, then what would be the meaning and nature of the latter? And, how valid are the conclusions about the ecosystem functioning obtained from them?. For the communities studied here, the low persistence of the consumer-resource interactions that constitute these networks has been highlighted ^17^. By combining this low interaction persistence with the high variability in degree distribution, time-aggregated networks likely represent all potential configurations and energy paths of the community as well as the ecological redundancy of species (nodes) and interactions (links). Time-aggregated networks, at least in our case, seem to represent more a map of potentialities, an attractor of the community dynamics, than a time-invariant representation of the ecological community as a natural unit. Novel perspectives, like those from multilayer network theory, offer analytical opportunities to deal with the challenge of properly describing community dynamics ^26^.

## Materials and Methods

### Biological data

We analyzed a long-term dataset of consumer-resource interactions obtained from macrobenthic intertidal rocky-shore communities in northern Chile. Four localities were sampled seasonally along ca.1,000 km of the northern coast of Chile: Río Seco (RS, 21.00°S, 70.17°W), Caleta Constitución (CC, 23.42°S, 70.59°W), Caleta Angosta (CA, 28.26°S, 71.38°W) and Lagunillas (LA, 30.10°S, 71.38°W). The sampling scheme yielded seasonal samples for each site, spanning the austral spring of 2004 and the austral summer of 2007. Within each site, 80% of the sampled individuals were identified at the species level, and trophic linkages were recorded from the analysis of stomach contents of each species. From these analyses, over 1,000 interactions in total were recorded ^27, 28^.

### Time-aggregated and seasonal networks

We constructed two types of networks from the biological data, namely seasonal and time-aggregated networks. Each seasonal network was constructed with all interactions observed in that given season, year, and site (n = 40). Time-aggregated networks were constructed by combining all of the observed interactions from the ten seasonal samplings at each site (n = 4). For both types of networks, the consumer-resource matrix contained binary values of 0 and 1, where 1 indicates that the interaction was observed and 0 indicates otherwise.

### Degree distribution models

For both types of networks, we quantified the degree distribution, which describes the frequency distribution of links per species. We evaluated the in-degree (number of prey consumed by each predator) and the out-degree (number of predators attacking each prey) by fitting two candidate models: the discrete version of a generalized beta distribution (DGBD) and an exponential distribution. The DGBD has the form:

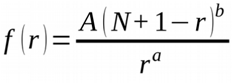

where r is the rank value of the node in and out-degrees, N is the maximum possible rank value, A is a normalization constant, and *a* and *b* are exponents. The exponent *a* is related to the “left to right” tail and the exponent *b* is related to the “right to left” tail of the dataset; thus, different combinations of values of *a* and *b* correspond to different shapes of the curve ^25^. In particular, parameter *a* is an approximated equivalent of the exponent α of the power law formula commonly used to analyze the degree distribution of other networks ^18^, and for this reason we focused our analysis on this parameter. The rank exponential distribution it is defined by:

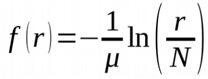

where r is the rank value of the node in and out-degrees, N is the maximum possible value of the rank, and μ is the decay exponent ^26^.

To determine the time at which the aggregation of seasonal networks converged to the estimated parameter values of the time aggregated networks, we sequentially added each seasonal matrix and fitted the DGBD and exponential functions to the observed degree distribution. We calculated the time point at which the aggregation of seasonal networks produced parameters similar to those of the aggregated networks (estimation ± standard error). This procedure is analogous to methods used to estimate species rarefaction curves. All analyses were carried out in the R environment ^29^ using the network ^30^ and igraph ^31^packages.

## Acknowledgements

We thank financial support from CONICYT grant N° 21140959 to DNL, FONDECYT 1040425 to PAC, FONDECYT 1141037 and FONDAP IDEAL 15150003 to NV, and Center of Applied Ecology & Sustainability (CAPES) FB 0002, FONDECYT 1160370 and FIA-PYT-2016-0203 to SAE. Dr. Fabio Labra and Dr. Sebastian Abades provided insightful comments that improved a previous version of this manuscript.

## Author contributions

All co-authors have substantially contributed to conception and design, acquisition of data, and analysis and interpretation of data; drafted or critically revised the article for important intellectual content Lopez, Daniela N. wrote the first draft of the manuscript and analyzed data, Camus, Patricio A. facilitated the data set, Valdivia, Nelson analyzed output data & Estay, Sergio A. performed statistical analysis and analyzed output data and all authors contributed substantially to revisions.

## References

1. Olesen, J. M. et al. From Broadstone to Zackenberg: Space, time and hierarchies in ecological networks. Adv. Ecol. Res. 42, 1 (2010).

2. Blonder, B., Wey, T. W., Dornhaus, A., James, R. & Sih, A. Temporal dynamics and network analysis. Methods Ecol. Evol. 3, 958–972 (2012).

3. Perra, N., Gonçalves, B., Pastor-Satorras, R. & Vespignani, A. Activity driven modeling of time varying networks. arXiv Prepr. arXiv1203.5351 (2012).

4. Holme, P. & Saramäki, J. Temporal networks. Phys. Rep. 519, 97–125 (2012).

5. Scholtes, I., Wider, N. & Garas, A. Higher-order aggregate networks in the analysis of temporal networks: path structures and centralities. Eur. Phys. J. B 89, 1–15 (2016).

6. Boccaletti, S., Latora, V., Moreno, Y., Chavez, M. & Hwang, D. Complex networks: Structure and dynamics. Phys. Rep. 424, 175–308 (2006).

7. Albert, R. Statistical mechanics of complex networks. Rev. Mod. Phys. 74, (2002).

8. Digel, C., Riede, J. O. & Brose, U. Body sizes, cumulative and allometric degree distributions across natural food webs. Oikos 120, 503–509 (2011).

9. Dunne, J. A., Williams, R. J. & Martinez, N. D. Small networks but not small worlds: unique aspects of food web structure. Proc. Nat. Acad. Sci (2002).

10. Dunne, J. A., Williams, R. J. & Martinez, N. D. Network Structure and Robustness of Marine Food Webs. (2003).

11. Estrada, E. Food webs robustness to biodiversity loss: the roles of connectance, expansibility and degree distribution. J. Theor. Biol. 244, 296–307 (2007).

12. Pascual & Dunne. Ecological Networks: Linking Structure to Dynamics in Food Webs. 2005, (2005).

13. Lurgi, M., López, B. C. & Montoya, J. M. Climate change impacts on body size and food web structure on mountain ecosystems. Philos. Trans. R. Soc. London B Biol. Sci. 367, 3050–3057 (2012).

14. Newman, M. E. J. The structure and function of complex networks. SIAM Rev. 45, 167–256 (2003).

15. Liu, J., Dang, Y., Wang, Z. & Zhou, T. Relationship between the in-degree and out-degree of WWW. Phys. A Stat. Mech. its Appl. 371, 861–869 (2006).

16. Xu, Y., Liu, P., Li, X. & Ren, W. Discovering the influences of complex network effects on recovering large scale multiagent systems. ScientificWorldJournal. 407639 (2014). doi:10.1155/2014/407639

17. Lopez, D. N., Camus, P. A., Valdivia, N. & Estay, S. A. High temporal variability in the occurrence of consumer-resource interactions in ecological networks. Oikos:DOI10.1111/oik.04285 (2017).

18. Blüthgen, N., Menzel, F. & Blüthgen, N. Measuring specialization in species interaction networks. BMC Ecol. 6, 9 (2006).

19. Holt, R. D. Predation, apparent competition, and the structure of prey communities. Theor. Popul. Biol. 12, 197–229 (1977).

20. Jeffries, M. J. & Lawton, J. H. Enemy free space and the structure of ecological communities. Biol. J. Linn. Soc. 23, 269–286 (1984).

21. Albert, R. The robustness of complex networks. in APS March Meeting Abstracts 1, 3002 (2002).

22. Paine, R. T. Food web complexity and species diversity. Am. Nat. 65–75 (1966).

23. Luo, J. & Whitney, D. E. Asymmetry in in-degree and out-degree distributions of large-scale industrial networks. arXiv Prepr. arXiv1507.04507 (2015).

24. Holt, R. D. in Food webs 313–323 (Springer, 1996).

25. Proulx, S. R., Promislow, D. E. L. & Phillips, P. C. Network thinking in ecology and evolution. Trends Ecol. Evol. 20, 345–353 (2005).

26. Pilosof, S., Porter, M. A., Pascual, M. & Kéfi, S. The multilayer nature of ecological networks. Nat. Ecol. Evol. 1, 101 (2017).

27. Camus, P. & Daroch, K. Potential for omnivory and apparent intraguild predation in rocky intertidal herbivore assemblages from northern Chile. Mar. Ecol. Prog. Ser. 361, 35–45 (2008).

28. Camus, P. A., Arancibia, P. A. & Ávila-Thieme, I. A trophic characterization of intertidal consumers on Chilean rocky shores. Rev. Biol. Mar. Oceanogr. 48, 431–450 (2013).

29. RCore, T. R: A language and environment for statistical computing. R Foundation for Statistical Computing, Vienna, Austria. *Online http://www.R-project.org* (2013).

30. Butts, C. T. network: a Package for Managing Relational Data in R. J. Stat. Softw. 24, 1–36 (2008).

31. Csardi, G. & Nepusz, T. The igraph software package for complex network research. InterJournal, Complex Syst. 1695, 1–9 (2006).

